# XpBrew and PanXpresso - automatic RNA-seq processing workflow and comprehensive collection of gene expression data

**DOI:** 10.64898/2026.08.27.747620

**Authors:** Shakunthala Natarajan, Claudia Sterling, Nancy Choudhary, Najnin Khatun, Mia-Sophie Brieske, Hannah Elisa Busch, Boas Pucker

## Abstract

Rapid developments in sequencing technologies have reduced the costs of transcriptomic experiments and resulted in a plethora of publicly available RNA-seq datasets. This is a valuable resource that can be harnessed to obtain novel biological insights through data upcycling. In this wake, we introduce XpBrew, an end-to-end Python workflow that was applied to generate PanXpresso, a comprehensive collection of gene expression datasets covering the taxonomic breadth of plants, animals, fungi, bacteria and archaea. XpBrew (https://github.com/PuckerLab/XpBrew) and PanXpresso (https://doi.org/10.60507/FK2/OBIGQH) are freely available.

## Introduction

Sequencing technologies, including those for transcriptomics have been subjected to a number of transition stages and have witnessed rapid advancements leading to substantially reduced costs for experiments^1^. Beginning with expressed sequence tags (ESTs), and microarrays, the gene expression analysis domain has witnessed the emergence of RNA-seq analysis and finally single-cell and spatial transcriptomics technologies more recently^2^. RNA-seq has widely replaced microarrays for gene expression quantification. This gene expression analysis landscape in particular has been shaped by short reads (Illumina); with long read methods offered by PacBio and Oxford Nanopore Technologies emerging later^3^. RNA-seq is essentially the sequencing of complementary Deoxyribonucleic Acid (cDNA) fragments and provides a clear snapshot of the dynamic transcript levels in an organism at a given time point under certain conditions. A transcriptome, as the name suggests, is a collection of transcripts in an organism’s tissue at a given time point under specific conditions. Key applications of RNA-seq include: (1) *de novo* transcriptome assembly, (2) generation of RNA-seq hints for structural annotation, and (3) gene expression quantification^3^. These applications are further followed-up by differential gene expression analysis and ultimately gene function annotation while also helping understand the regulatory triggers that can be inferred only via expression analyses^4^. RNA-seq, based on short read technologies, provides high throughput at low costs, which is ideal for transcriptomic studies requiring large numbers of reads to infer transcript abundances. Long read technologies like IsoSeq and direct RNA sequencing are better suited for isoform quantification, but high associated costs and resource requirements outweigh their utility in most applications^5^.

The RNA-seq reads generated in studies employing these different methods are deposited primarily in the Sequence Read Archive (SRA) hosted by the National Center for Biotechnology Information (NCBI)^6^, the European Nucleotide Archive (ENA)^7^, or DDBJ Sequence Read Archive (DRA) hosted by DNA Data Bank of Japan (DDBJ)^8^. Data submitted to ENA or Gene Expression Omnibus (GEO) are reflected in NCBI SRA, making it ‘the’ repository for retrieving RNA-seq read data for expression analysis. The rise of transcriptome technologies, combined with the rapidly increasing RNA-seq read dataset numbers, has also expanded the utility and application of RNA-seq data in general^9^. While the raw RNA-seq read data are hosted in some of the aforementioned public repositories and have been used in independent studies for expression analyses, there is still a dearth of organized repositories or data collections that can facilitate easy access to expression data that can further facilitate immediate downstream analyses and can be accessed by biologists with varying levels of computational expertise. PlantExp is a web platform that hosts expression data and alternative splicing profiles of 85 plant species across 24 plant orders with rich visualization options^10^. Following this, Plant Expression Omnibus (PEO) was released as a web application that hosts processed gene expression datasets of over 100 plant species. It also provides functional annotation for each gene, allowing for comparative genomics across the kingdom^11^. FungiExpresZ is a fungal gene expression database encompassing processed RNA-seq data from 23 different fungal species, accompanied by downloadable gene ontology (GO) data and several plotting and visualization options. This webpage is accompanied by a web tool also available as an R shiny package to process fungal expression data^12^. Later in 2024, a comprehensive gene expression collection of over 200 plant species was released as a data publication that provided the processed expression data files, coding sequence (CDS) files, and functional annotation files^13^. Some of the aforementioned repositories are either not accessible or do not have updated expression datasets. Given the rate at which repositories like SRA are teeming with transcriptomic datasets, availability of an updated expression collection that encompasses the new datasets will enable biological analyses at a fine-grained level and depth. Such large-scale expression data can facilitate a large number of studies pertaining to gene expression distribution analysis and gene function elucidation. For instance, biosynthetic pathway discovery is a key research area that benefits from the availability of large-scale transcriptomic datasets for coexpression analyses. For example, such readily available expression datasets can facilitate co-expression analysis across species borders by CoExpPhylo^14^. Spatial transcriptomics can also be performed with the availability of relevant metadata. Further applications include sample classification and identification of marker genes associated with specific conditions and/or phenotype(s) ^11,15^.

In this study, we introduce XpBrew, a Python-based pipeline to facilitate gene expression data processing. It allows for a reproducible generation of these expression datasets and also makes future updates convenient. XpBrew was used to generate PanXpresso - a massive collection of gene expression data across the domains of plants, animals, fungi, archaea, and bacteria. To the best of our knowledge, this is the first comprehensive expression data collection encompassing all major domains of life. The data collection is organized domain-wise and within each domain, species-wise. Every species’ processed gene expression data in transcripts per million (TPM), along with the coding sequence (CDS) file used for transcript abundance quantification and a derived polypeptide sequence set (PEP), have been provided.

## Results

### XpBrew enables effective RNA-seq data processing and convenient count table updating

Retrieving and processing large numbers of RNA-seq datasets can be a time-consuming endeavor. To automate the process and to make it reproducible, the individual steps have been integrated into XpBrew (Figure 1). This workflow covers the data retrieval in the most effective way, the automatic analysis with kallisto to obtain gene expression values, and downstream filtering to ensure that only valid RNA-seq datasets are included in the final collection. XpBrew also supports updating existing count tables with newly released datasets by only processing these additional datasets thus contributing to sustainability.

**Figure 1.**
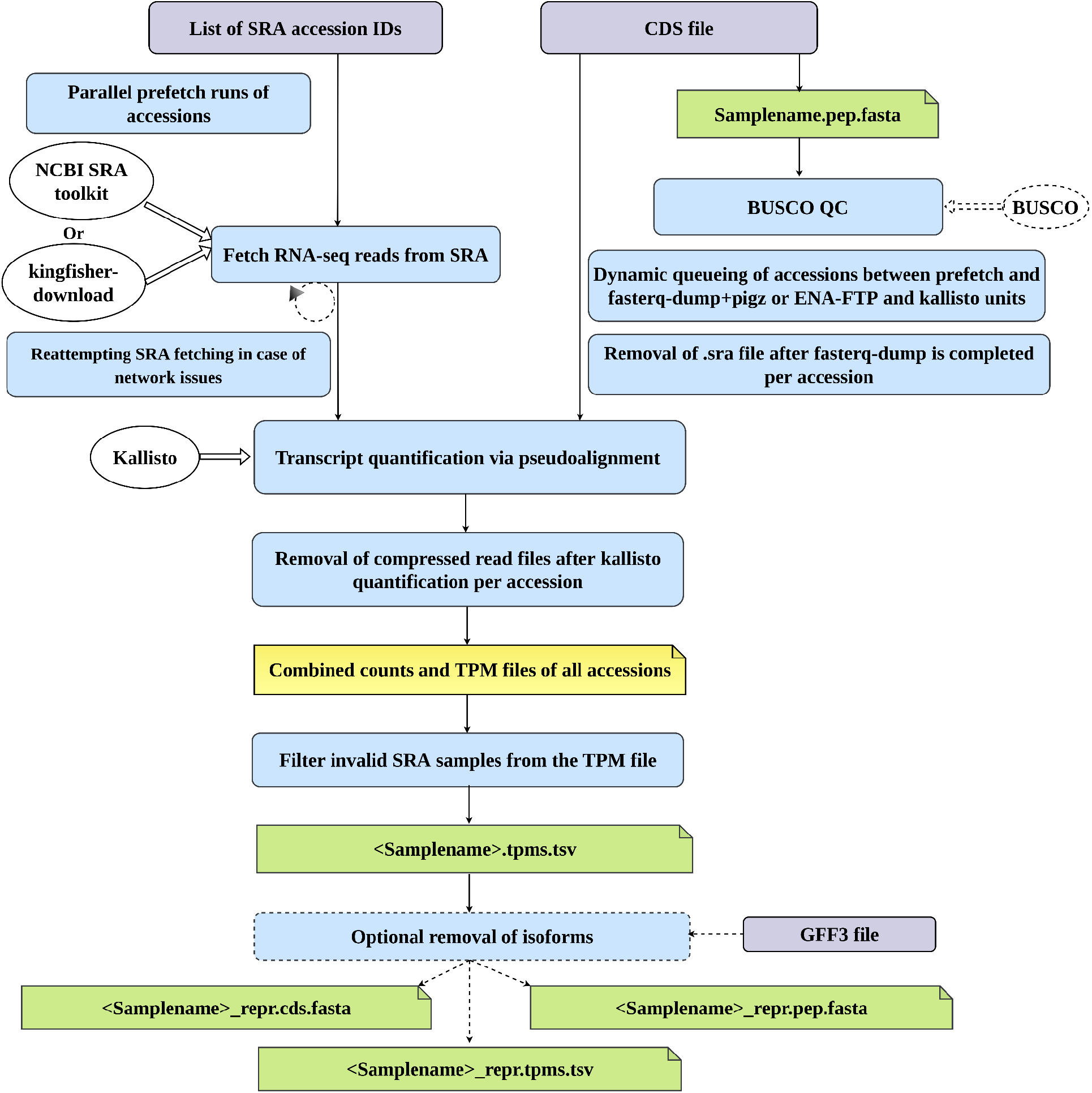
Schematic representation of XpBrew pipeline workflow. Purple boxes indicate user-provided input files. Blue boxes indicate the pipeline steps. Yellow and green page boxes represent the intermediate and final output files, respectively. External tools are shown by the white ovals. Representations with dotted outlines indicate optional steps.

The runtime is heavily dependent on the available internet connection, the computational environment available for executing XpBrew, the complexity of the reference, and the size of the RNA-seq dataset. In our experience, several hundred RNA-seq datasets of average size can be processed on 32 CPUs per day (de.NBI cloud).

### PanXpresso makes gene expression information accessible

The expression analysis with XpBrew resulted in count tables comprising transcript per million (TPM) values of 424 species belonging to the domains plantae, fungi, animalia, bacteria, and archaea (Table 1). Table 1 lists the order, family, species-wise coverage of the expression datasets. The taxonomic classification is based on information provided by the NCBI. One RNA-seq sample refers to an individual run provided by SRA/ENA. Only samples that passed all quality filters were included. A total of 5,07,331 RNA-seq data sets have been successfully processed and passed all quality filters (Supplementary File 1).

**Table 1.** Summary of gene expression data in PanXpresso.

| **Domain** | **Number of orders represented** | **Number of families represented** | **Number of species represented** | **Number of RNA-seq samples processed** |
| --- | --- | --- | --- | --- |
| Plantae | 41 | 83 | 342 | 325703 |
| Animalia | 19 | 25 | 28 | 102066 |
| Fungi | 11 | 13 | 21 | 39833 |
| Bacteria | 12 | 17 | 24 | 38897 |
| Archaea | 6 | 7 | 9 | 832 |

PanXpresso covers 75 angiosperm families and 8 non-angiosperm families (Figure 2), which enables systematic analyses across the taxonomic diversity of plants. The inclusion of fungal, animal, bacterial and archaeal expression datasets in PanXpresso enables cross-domain expression analyses (Figure 3). The completeness of the coding sequence collection used as reference for transcript quantification in this study was assessed with BUSCO (Supplementary File 1, Supplementary File 3). Priority was given to species with a high number of available RNA-seq datasets, which often resulted in the inclusion of well-studied model organisms (Figure 2, Figure 3) (Supplementary File 1). A detailed list of species, their taxonomy classification, BUSCO completeness and the number of SRA samples processed per-species is summarized in Supplementary File 1.

**Figure 2.**
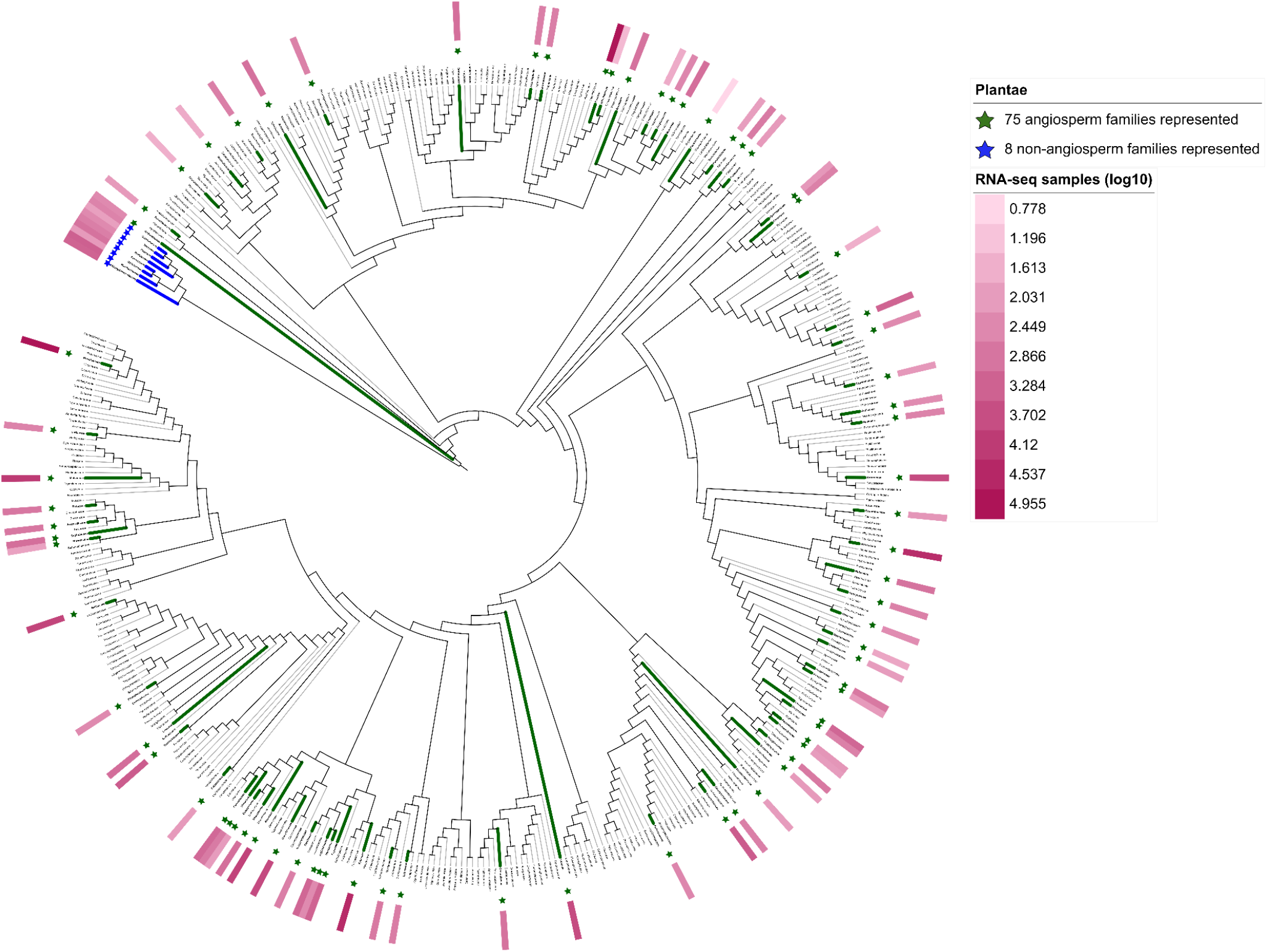
Phylogenetic coverage of plant families in PanXpresso. Plant families covered by gene expression data in PanXpresso are indicated with asterisks and highlighted branch colors. The number of RNA-seq samples per plant family are represented by the heat map. The RNA-seq sample numbers are log-transformed to the base of 10. The phylogenetic tree has been adapted and modified from Zuntini *et al*., 2024^16^

**Figure 3.**
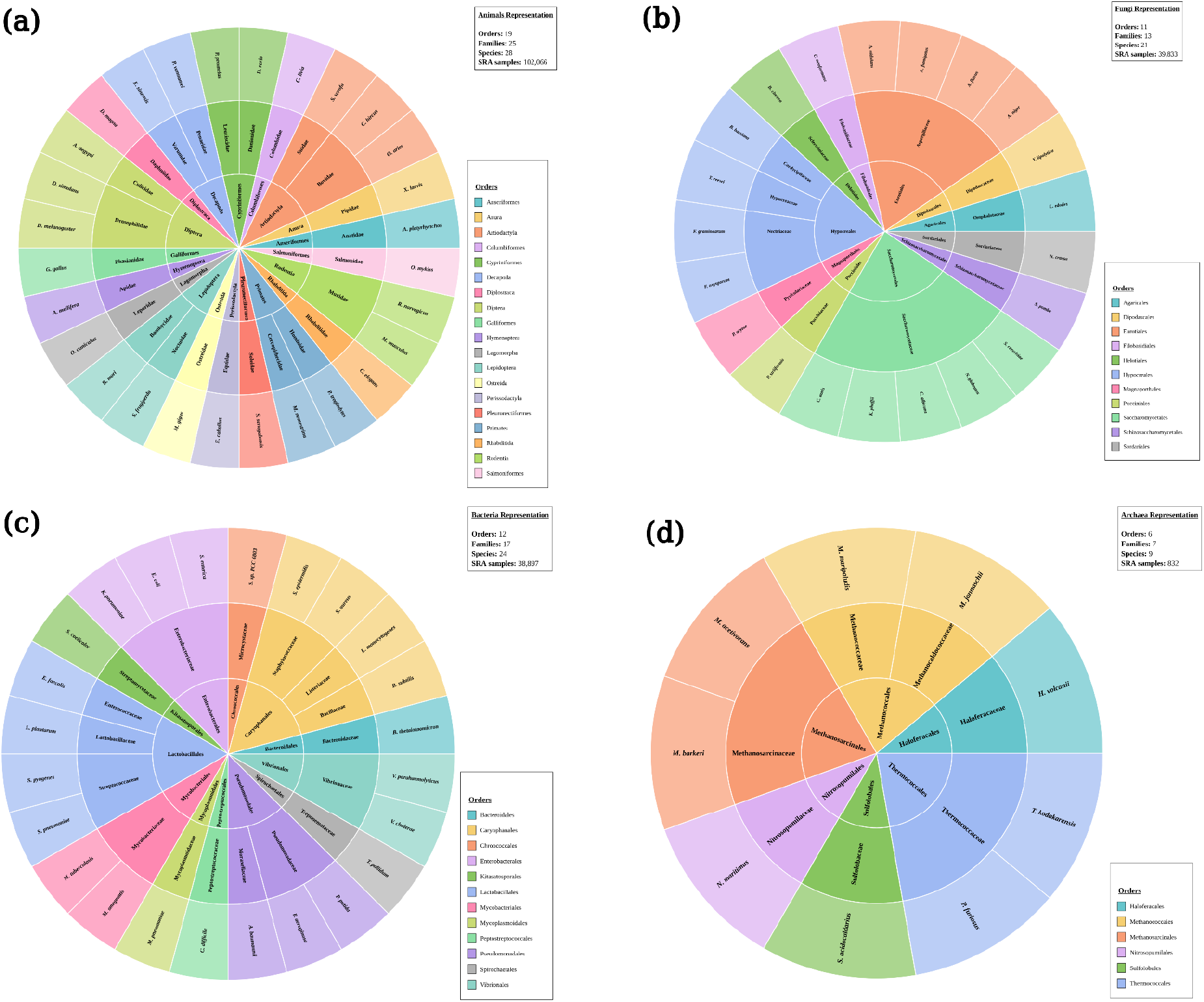
Sunburst plots depicting the diversity of non-plant domains (a) animals (b) fungi represented (c) bacteria (d) archaea in PanXpresso. Taxonomic information provided by the NCBI has been used for the classification in this figure.

## Discussion

This study resulted in the development of a comprehensive gene expression data collection of organisms from major representative orders across the domains of life. According to the hierarchical reports of the Integrated Taxonomic Information System (ITIS), there are approximately 969 plant families, 6450 animal families, 630 fungi families, 337 bacterial families, and 28 archaea families reported on Earth.

Among plants, angiosperms, the largest group of flowering plants, are well studied. The angiosperms family tree from Zuntini et al., 2024 represents 424 angiosperm families that translate to ~86% of the total known angiosperm diversity. PanXpresso covers 75 angiosperm families that is nearly 20% of the known angiosperm diversity and 8 non-angiosperm families, representing around 9% of the known plant families. The quality of the annotations underlying the expression collection is evident from Supplementary File 3 wherein majority of the datasets across all the domains show a BUSCO completeness greater than 90%, making it a diverse and high quality expression data collection. From Table 1, it is inferred that roughly 1% (animalia), 2% (fungi), 5% (bacteria), and 28% (archaea) of the families across domains are represented in the expression collection. These numbers indicate the coverage width of the expression data collection we present. As such, this collection offers good coverage for the domains plantae and archaea and provides datasets of well-studied species belonging to major families from the other domains, making it a comprehensive expression collection across the tree of life in relative comparison to the existing expression data repositories. Nevertheless, these statistics also indicate that future updates must focus on the underrepresented families to increase the diversity representation. There is an imbalance in datasets generated for model and non-model species across all the domains. The heatmap depicted as a part of the phylogenetic tree (Figure 2) provides a visual cue for identifying expression hotspot families harbouring plant species that have the most and least RNA-seq samples (indicated by the colour density). Sample proportioned sunburst plots for non-plant domains (Supplementary File 2) also provide information on families with most and least RNA-seq datasets. The potential for obtaining an even larger-scale transcriptomic data collection showing a higher diversity of the species is thus limited by the dataset skew leaning towards model species. Therefore more experimental work covering a wider diversity of species coupled with regular processing and updates of expression collection repositories can help in the creation of state-of-the-art consortium-scale expression collection that can facilitate downstream biological analyses at an unprecedented depth and rate^2,17^.

That said, it is also important to note that only paired-end RNA-seq datasets have been used to obtain this gene expression collection, as specified before. Further, the data processing pipeline XpBrew uses kallisto at its core, which can only compute average fragment lengths for paired-end read datasets automatically^18,19^. Given that no metadata about the library characteristics are available for all single-end datasets, these cannot be processed with high reliability. Moreover, the XpBrew pipeline employs the pseudo-alignment approach via kallisto for gene expression quantification, helping expedite the gene expression collection process while ensuring accuracy equivalent to mapping-based methods^18^. With the input requirement of only CDS (reference) and SRA lists from the user, the pipeline simplifies RNA-seq data extraction and processing steps, eliminating intermediate processing from the user’s end. With options to choose fetching via ENA-FTP (by integrating kingfisher-download^20^) or SRA toolkit^21^ with an optional fall-back to the latter, and the ability to restart after interruptions, the pipeline is a flexible and lightweight tool for expression analysis. The combination of our easily downloadable datasets along with the pipeline tool to facilitate such data collections in the future makes our data collection sustainable and modular with an easy option to update in the next coming years.

We provide an accessible data collection with a digital object identifier (https://doi.org/10.60507/FK2/OBIGQHI). The datasets are hosted in bonndata, the institutional research data repository of the University of Bonn, ensuring its continued persistence.Furthermore, disseminating the datasets as gene expression TPM files that can be readily processed for downstream analyses combats the global inequality pertaining to the availability of sufficient computational resources needed to process such massive datasets. This in turn helps democratize biological big data and promote inclusivity in science. Furthermore, large carbon footprints are associated with these significant computational resources needed for such big data endeavours^22^. Through this data collection, we mitigate the need for everyone to repetitively run such energy intensive operations. Lastly, given the impacts of political instability on data availability and maintenance in major research hubs^23^, it is important we generate and maintain important data resources in Europe. Our massive expression data collection - ‘PanXpresso’ is a definitive step in this direction. PanXpresso conforms to the FAIR principles of being findable, accessible, reusable and interoperable, making it a valuable European data resource for biological explorations and analyses by the scientific community at large.

## Methods

### Data collection

First, the major genome sequence repositories - NCBI^21^, Genome Warehouse^24^, and Phytozome^25^, were explored to obtain a comprehensive list of plants, fungi, bacteria, and animals, representative of the major orders in each kingdom. The genome sequence and corresponding annotation were retrieved in FASTA and GFF format, respectively. The CDS files were derived from the annotation with the Python script get_cds_from_assembly.py (v0.2) to ensure the use of clean sequence headers for all downstream analysis steps. Sequence IDs in the CDS files were ensured to match the IDs in the corresponding GFF files to later facilitate isoform removal based on the GFF file. RNA-seq datasets for each of the species included in the data collection were obtained from the SRA. The following filters were applied in the SRA sample selection process: (1) the specific species name was chosen in the taxonomy section, (2) RNA was set as the source, (3) paired-end library layout, and (4) FASTQ file types from Illumina and Beijing Genomics Institute (BGISEQ) sequencing platforms. The SRA accessions list file and the corresponding CDS and GFF file of each species were provided to XpBrew.py (v0.3.0) for retrieval and processing of the datasets. The GFF file was included to obtain a primary isoform retained version of the expression, CDS and PEP files along with the complete datasets including all isoforms. A primary isoform in the scope of this study is defined as the transcript with the longest coding sequence amongst all the transcripts of a gene.

### Gene expression analysis

XpBrew (Figure 1) starts with an optional step involving Benchmarking Universal Single Copy Orthologs (BUSCO) (v6.1.0)^26^ in the protein mode that uses the PEP file generated by in silico translation from the input CDS file. It then provides options for choosing the fetch method via kingfisher-download (v0.5.0)^20^ (currently only the ENA-FTP fetch method is enabled via kingfisher-download) or sratoolkit (v3.4.1)^21^ that uses prefetch (v3.4.1)^21^, fasterq-dump (v3.4.1)^21^, and pigz (v2.8)^27^ to obtain gzipped FASTQ files. Later, it employs kallisto (v0.52.0)^18^ for pseudo-alignment quantification of gene expression in transcripts per million (TPM) (https://github.com/bpucker/CoExp/kallisto_pipeline3.py). It includes a merge step to combine the count tables from the different samples (https://github.com/bpucker/CoExp/merge_kallisto_output3.py) and a pre-final filtration step. Samples with less than one million reads or samples where the percentage of reads assigned to the most abundant 100 transcripts fell outside the 10-80% range were filtered out as invalid samples (https://github.com/bpucker/CoExp/filter_RNAseq_samples.py). Additionally, it also provides options to merge the TPM expression file output produced in the current run with other TPM files that can be indicated via a config file, facilitating easy merge options for future updates and opportunities to process partial datasets in parallel on different systems. The pipeline employs a parallel prefetch. Afterwards, fasterq-dump and kallisto can effectively utilize all available cores for the data processing. The composite design is wrapped in a queue-fetch architecture wherein a prefetch-completed accession queues up for the serial block processing. The pipeline also checks for storage space availability to utilize the available disk space effectively, while avoiding exceeding the disk quota. Lastly, the pipeline also provides support for processing multiple references (multiple CDS files) concurrently in a single run, avoiding the need to fetch and download read data multiple times, thus making it more sustainable. Claude (model: Sonnet 4.6) was used to optimize the XpBrew pipeline.

### Summarizing and visualizing the dataset

Once all the datasets were collected, comprehensive checks were conducted to ensure correct header compatibility across the different output files per species. Checksums were also generated per species folder to ensure correct file identities and to facilitate reliable downloads. The quality of the datasets included in the collection, indicated by BUSCO completeness, was visualized with the help of radial wheel plots. The plant species included in the collection were classified into their respective families, and this classification was used to modify and annotate the family-level angiosperm phylogenetic tree file generated by Zuntini *et al*., 2024^28^. Since the collection included non-angiosperm plant species as well, the families of these plants were used to obtain a phylogenetic tree file using phyloT v2^29^ based on NCBI taxonomy with the following settings: internal nodes collapsed and no polytomy in the Newick format. This tree file was concatenated with the tree file from Zuntini *et al*., 2024 (ignoring branch lengths) to obtain a representative phylogenetic tree depicting the non-angiosperm families represented in PanXpresso along with the angiosperms. The other domains included in the collection - animals, fungi, bacteria, and archaea were visualized with the help of sunburst plots.

A comprehensive list of all the dataset sources has been consolidated in Supplementary File 1.

## Declarations

### Ethics approval and consent to participate

Not applicable

### Consent for publication

Not applicable

### Availability of data and materials

PanXpresso can be accessed via bonndata: https://doi.org/10.60507/FK2/OBIGQH. XpBrew software is freely available at: https://github.com/PuckerLab/XpBrew. Other scripts used for data processing in this study can be accessed at: https://github.com/ShakNat/Genomic_file_wrangler.

### Competing interests

The authors declare that they have no competing interests.

### Funding

Not applicable

### Authors’ contributions

BP designed the research project and supervised the work. SN and BP wrote the software for retrieving and processing the datasets. SN, CS, NC, NJ, MSB, HEB, and BP performed the data collection and bioinformatics analyses. All the authors interpreted the results and SN and BP wrote the manuscript. All the authors approved the final version of the manuscript and agreed to its submission.

## Acknowledgements

This work was supported by the de.NBI Cloud within the German Network for Bioinformatics Infrastructure (de.NBI) and ELIXIR-DE (Forschungszentrum Jülich and W-de.NBI-001, W-de.NBI-004, W-de.NBI-008, W-de.NBI-010, W-de.NBI-013, W-de.NBI-014, W-de.NBI-016, W-de.NBI-022). SN and NK were supported by the doctoral scholarship of the German Academic Exchange Service (DAAD). We thank all members of the Plant Biotechnology and Bioinformatics group for their support and feedback during the process.

## Notes

https://github.com/PuckerLab/XpBrew

https://doi.org/10.60507/FK2/OBIGQH

### Competing Interest Statement

The authors have declared no competing interest.

