## Supplementary figures and images for "XpBrew and PanXpresso - automatic RNA-seq processing workflow and comprehensive collection of gene expression data"

### Supplementary File 2

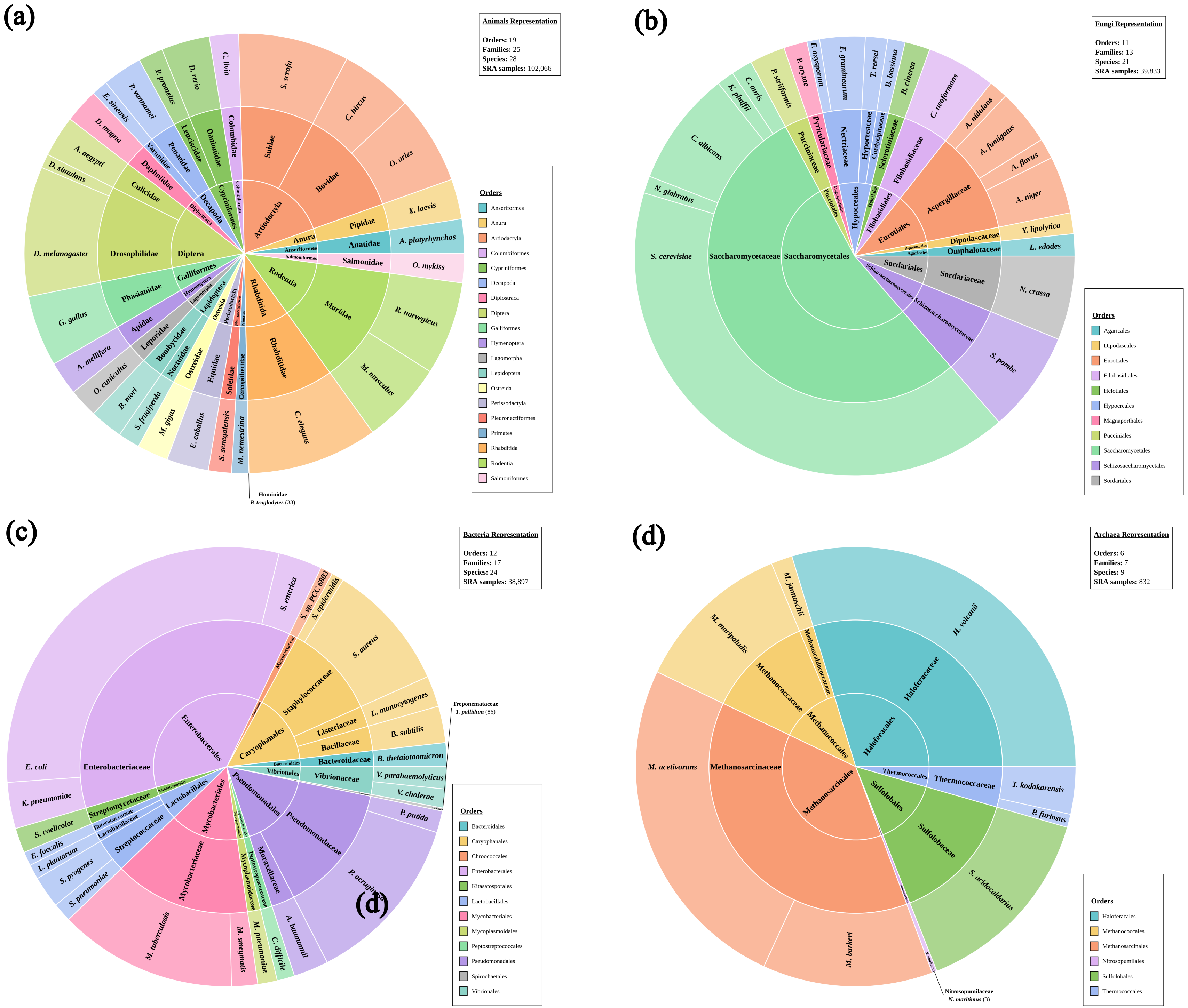


**RNA-seq sample proportional sunburst plots for (a) animals, (b) fungi, (c) bacteria, and (d) archaea**
