## Supplementary File 3 for "XpBrew and PanXpresso - automatic RNA-seq processing workflow and comprehensive collection of gene expression data"

(a)

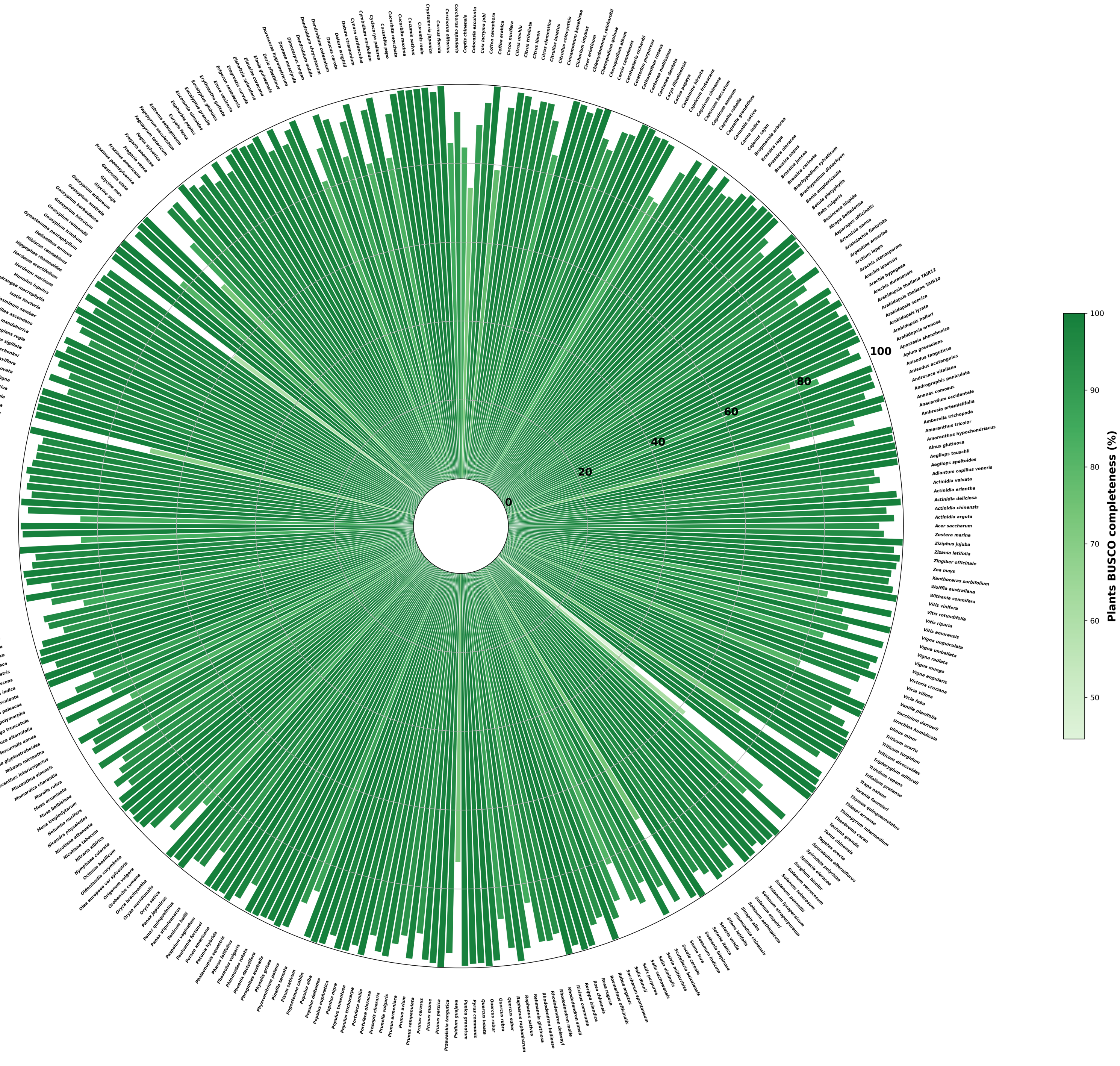

(b)

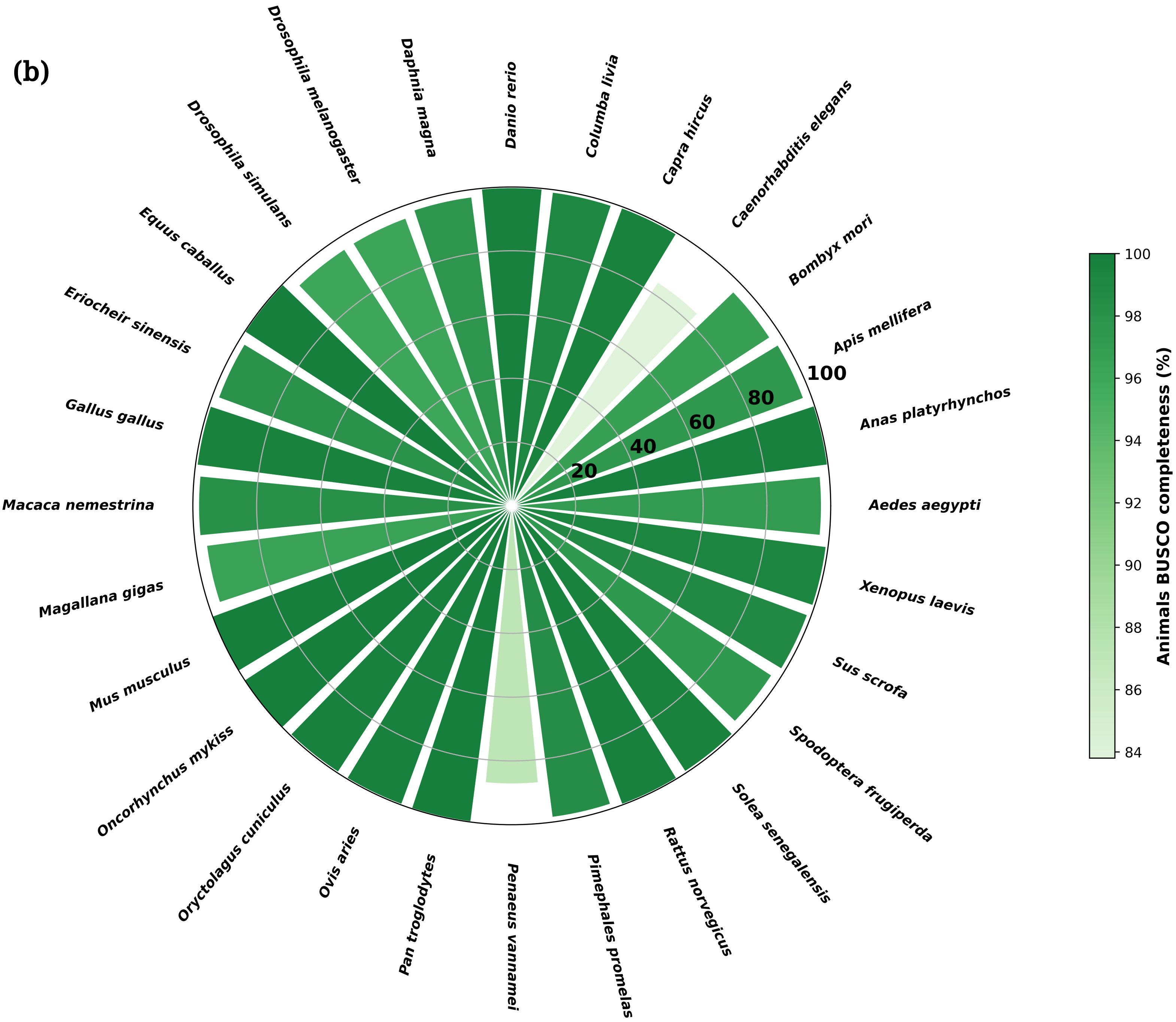

Danio rerio
Columba livia
Capra hircus
Caenorhabditis elegans
Bombyx mori
Apis mellifera
Anas platyrhynchos
Aedes aegypti
Xenopus laevis
Sus scrofa
Spodoptera frugiperda
Solea senegalensis
Rattus norvegicus
Pimephales promelas
Penaeus vannamei
Pan troglodytes
Ovis aries
Oryctolagus cuniculus
Oncorhynchus mykiss
Mus musculus
Magallana gigas
Macaca nemestrina
Gallus gallus
Eriocheir sinensis
Equus caballus
Drosophila simulans
Drosophila melanogaster
Daphnia magna
20
40
60
80
100
Animals BUSCO completeness (%)
84
86
88
90
92
94
96
98
100

(c)

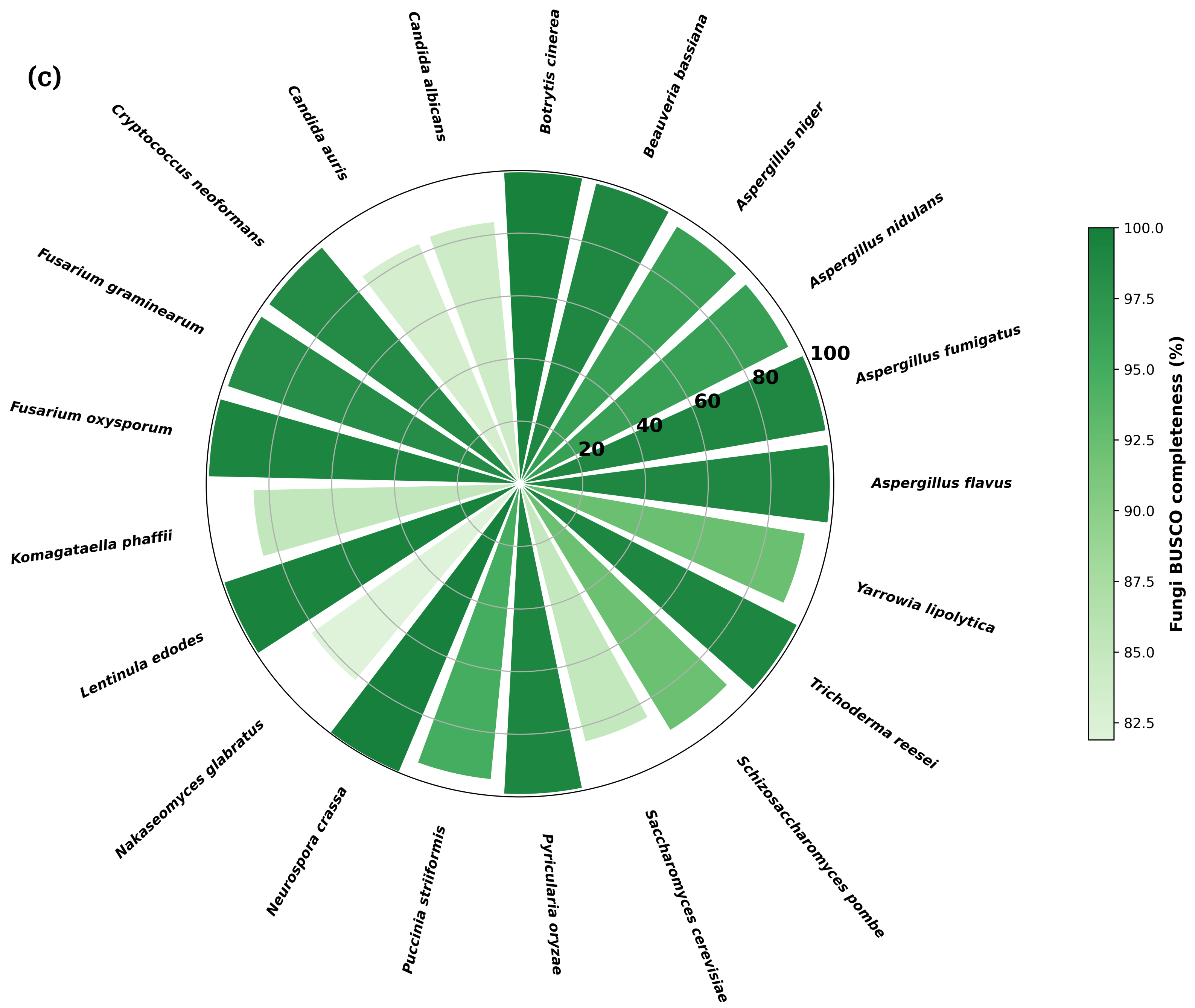
Botrytis cinerea
Beauveria bassiana
Aspergillus niger
Aspergillus nidulans
Aspergillus fumigatus
Aspergillus flavus
Yarrowia lipolytica
Trichoderma reesei
Schizosaccharomyces pombe
Saccharomyces cerevisiae
Pyricularia oryzae
Puccinia striiformis
Neurospora crassa
Nakaseomyces glabratus
Lentinula edodes
Komagataella phaffii
Fusarium oxysporum
Fusarium graminearum
Cryptococcus neoformans
Candida auris
Candida albicans
20
40
60
80
100
Fungi BUSCO completeness (%)
100.0
97.5
95.0
92.5
90.0
87.5
85.0
82.5

(d)

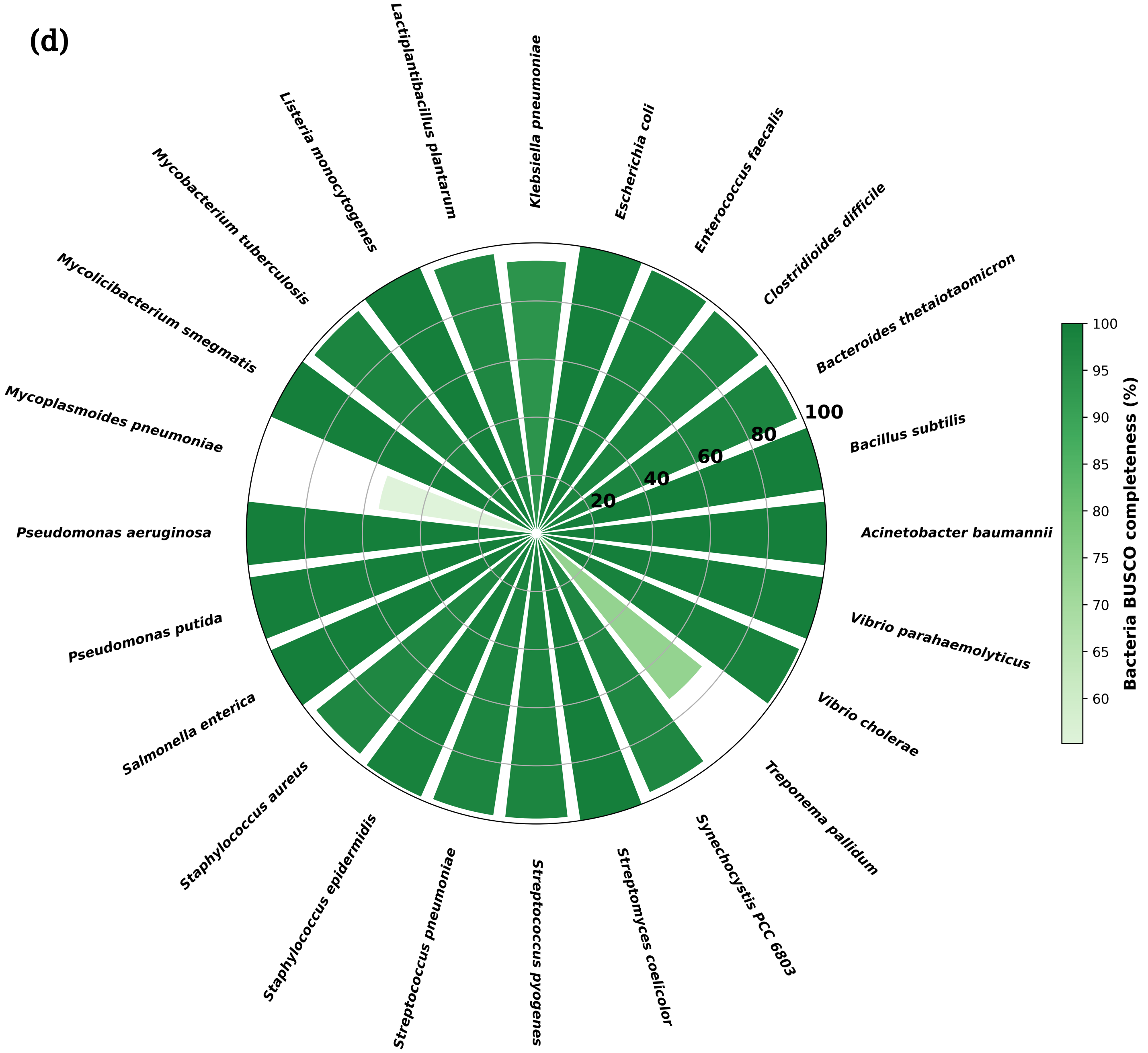
Klebsiella pneumoniae
Escherichia coli
Enterococcus faecalis
Clostridioides difficile
Bacteroides thetaiotaomicron
Bacillus subtilis
Acinetobacter baumannii
Vibrio parahaemolyticus
Vibrio cholerae
Treponema pallidum
Synechocystis PCC 6803
Streptomyces coelicolor
Streptococcus pyogenes
Streptococcus pneumoniae
Staphylococcus epidermidis
Staphylococcus aureus
Salmonella enterica
Pseudomonas putida
Pseudomonas aeruginosa
Mycoplasmoides pneumoniae
Mycolicibacterium smegmatis
Mycobacterium tuberculosis
Listeria monocytogenes
Lactiplantibacillus plantarum
20
40
60
80
100
Bacteria BUSCO completeness (%)
60
65
70
75
80
85
90
95
100

(e)

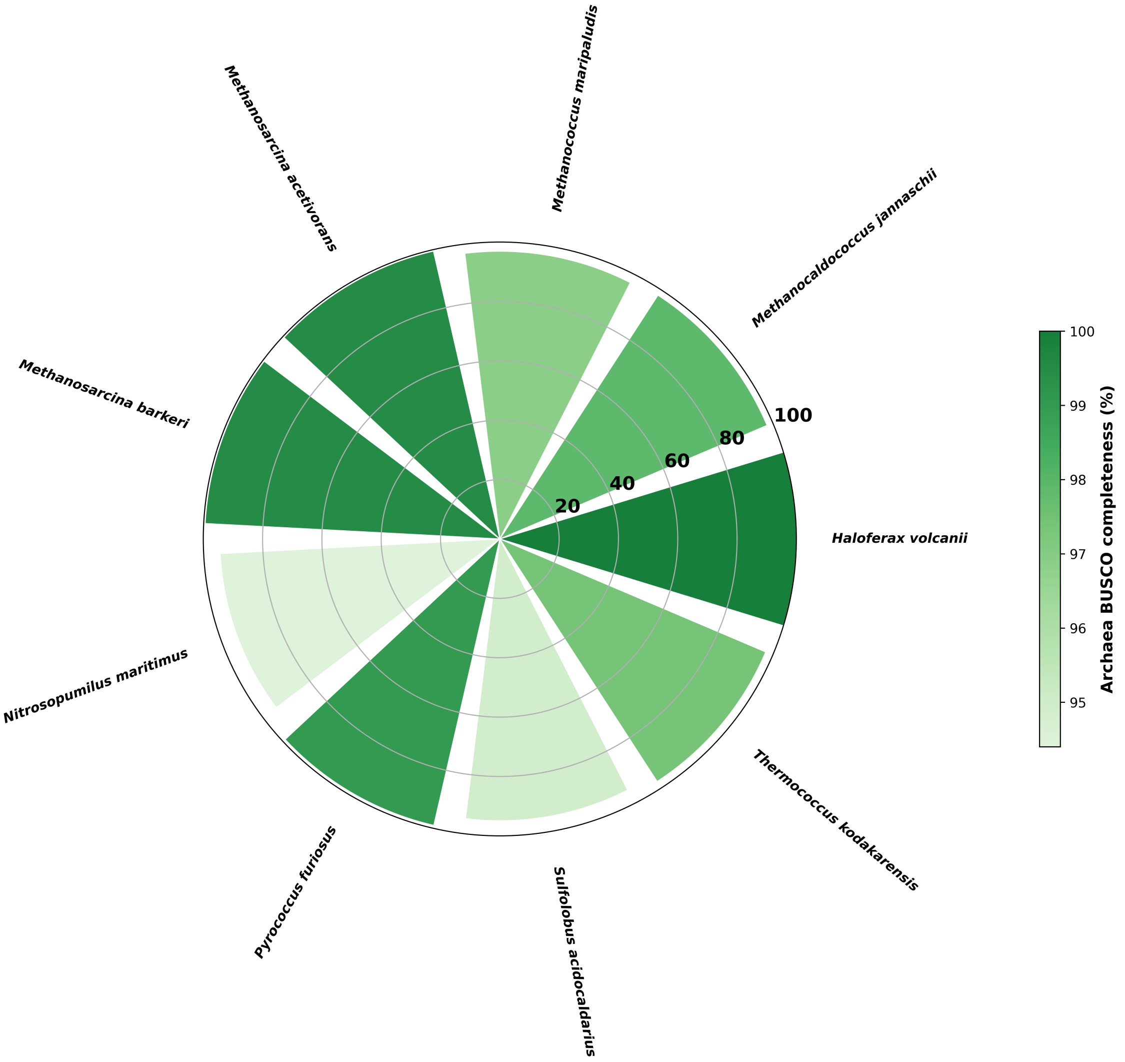


**BUSCO completeness plots based on polypeptide sequences of species across (a) plants (b) animals (c) fungi (d) bacteria and (e) archaea. The scores represented in the plots are based on polypeptide sequences of all transcripts. BUSCO completeness scores of isoform removed polypeptide sequences can be found in Supplementary File 1.**
